# BOLE: A Knowledge-Enhanced Multi-Agent Framework for Intelligent Genomic Breeding

**DOI:** 10.64898/2026.02.04.703673

**Authors:** Zishuai Wang, Chongxiao Liang, Xiaoai Zhang, Wenkang Wei, Kui Li

## Abstract

Genomic breeding has become increasingly data-intensive, yet the practical integration of heterogeneous bioinformatics tools into coherent analytical workflows remains a major bottleneck. To address this, we present BOLE, a knowledge-enhanced multi-agent AI framework for autonomous genomic breeding analysis. By integrating a structured script knowledge base with constrained large language model reasoning, BOLE translates high-level analytical intent into validated and executable workflows. Specialized agents collaboratively perform intent interpretation, task decomposition, dependency reasoning, workflow synthesis, and controlled code generation to assemble validated, executable workflows without predefined pipelines. BOLE supports core genomic breeding tasks, including genome-wide association studies, heritability estimation, germplasm evaluation, and genomic selection while preserving artifact-level provenance to ensure reproducibility. Implemented as a web service, BOLE enables end-to-end genomic breeding analysis from raw data to actionable results for non-expert users. This work establishes a generalizable paradigm in which knowledge-driven multi-agent AI bridges the gap between mature analytical methods and their real-world application in genomic breeding.

## 1. Introduction

The rapid accumulation of high-throughput genotyping and phenotype data has fundamentally transformed genomic breeding, enabling data-driven strategies such as genome-wide association studies (GWAS) and genomic selection (GS)^1,2^. These analytical approaches are now considered essential components of modern breeding programs aimed at improving economically important traits^3^. However, despite the methodological maturity of individual statistical models and software tools^4-6^, the practical execution of genomic breeding analyses remains a substantial bottleneck for many researchers and breeding practitioners. Real-world analyses require coordinated use of heterogeneous command-line tools, statistical environments, and custom scripts. Each analytical step—ranging from genotype quality control to downstream prediction modeling—imposes strict constraints on data formats, parameterization, and software dependencies. As a result, end users must possess not only domain knowledge in quantitative genetics but also advanced computational expertise to manually design, adapt, and maintain complex analysis workflows^7^. This reliance on expert-driven pipeline construction limits reproducibility, reduces accessibility for non-specialists, and hinders the scalable application of genomic breeding technologies.

Several workflow management platforms and web-based tools have been developed to mitigate these challenges by integrating commonly used genomic analysis software^8,9^. While such systems improve usability to some extent, most rely on predefined, static pipelines or manual configuration of analytical steps^10^. Consequently, users continue to shoulder the responsible for selecting appropriate methods, resolving data compatibility issues, and adapting workflows to diverse experimental designs. These limitations highlight a fundamental gap between the availability of powerful genomic analysis tools and the ability of end users to autonomously and reliably apply them in real-world breeding scenarios^11^.

Multi-agent systems (MAS) offer a promising paradigm for addressing this gap by enabling distributed reasoning, task decomposition, and autonomous decision-making in complex problem spaces^12,13^. In contrast to traditional workflow automation, MAS-based frameworks can explicitly model domain knowledge, reason over task dependencies, and dynamically construct executable processes based on user objectives and data characteristics^14^. Although MAS has been explored in domains such as robotics, manufacturing, and intelligent decision support^15^, its application to livestock genomic breeding analysis remains limited^16^.

In this study, we present BOLE, a knowledge-enhanced multi-agent AI framework for intelligent genomic breeding analysis. BOLE integrates a structured script knowledge base that formalizes analytical modules for GWAS, heritability estimation, germplasm evaluation, and genomic selection, including their input–output specifications and software dependencies. Through the collaboration of specialized agents for user intent interpretation, workflow planning, parameter harmonization, and script execution, the system dynamically assembles end-to-end analysis pipelines without requiring manual workflow design. To enable robust semantic interpretation and adaptive planning, BOLE leverages large language model (LLM)–driven reasoning. Crucially, all LLM outputs are constrained and checked against the formalized knowledge base and validation rules, and any auxiliary code synthesis is subject to schema conformance and automated verification before execution. We implement BOLE as a web-deployed platform that provides a lightweight interaction layer for submitting data and high-level analytical intents while delegating reasoning, planning, and execution to the multi-agent backend. By bridging expert knowledge and automated orchestration, BOLE aims to bridge the gap between methodological maturity and practical deployment in genomic breeding, offering a generalizable paradigm for intelligent, scalable decision support in data-intensive breeding programs.

## 2. Materials and Methods

### 2.1. System Architecture and Multi-Agent Framework of BOLE

The system is implemented as a web-based platform utilizing a MAS architecture to manage complex genomic workflows (Figure 1). The backend is developed using the FastAPI framework^17^, which serves as the orchestration hub for specialized agents. The frontend is constructed with React and Vite, providing a terminal-based interface for real-time execution monitoring and a natural language interaction environment. The MAS architecture comprises four primary functional agents: Interaction Agent: Interacts with users to interpret high-level analysis objectives and transform natural-language inputs into structured analytical intents. Plan Agent: Decomposes user intents into a sequence of analytical tasks and determines feasible execution paths based on data availability and task dependencies. Pipeline Assembly Agent: Automatically resolves data format mismatches by inserting intermediate transformation steps when necessary, ensuring compatibility between successive tasks. Code Generation Agent: Dynamically synthesizes self-sufficient, executable scripts in R, Python, or Bash to fulfill custom analytical requirements and visualization tasks not covered by predefined pipelines. Through agent collaboration, BOLE achieves distributed reasoning and task coordination, allowing the system to autonomously adapt workflows to different datasets and analysis goals.

**Figure 1.**
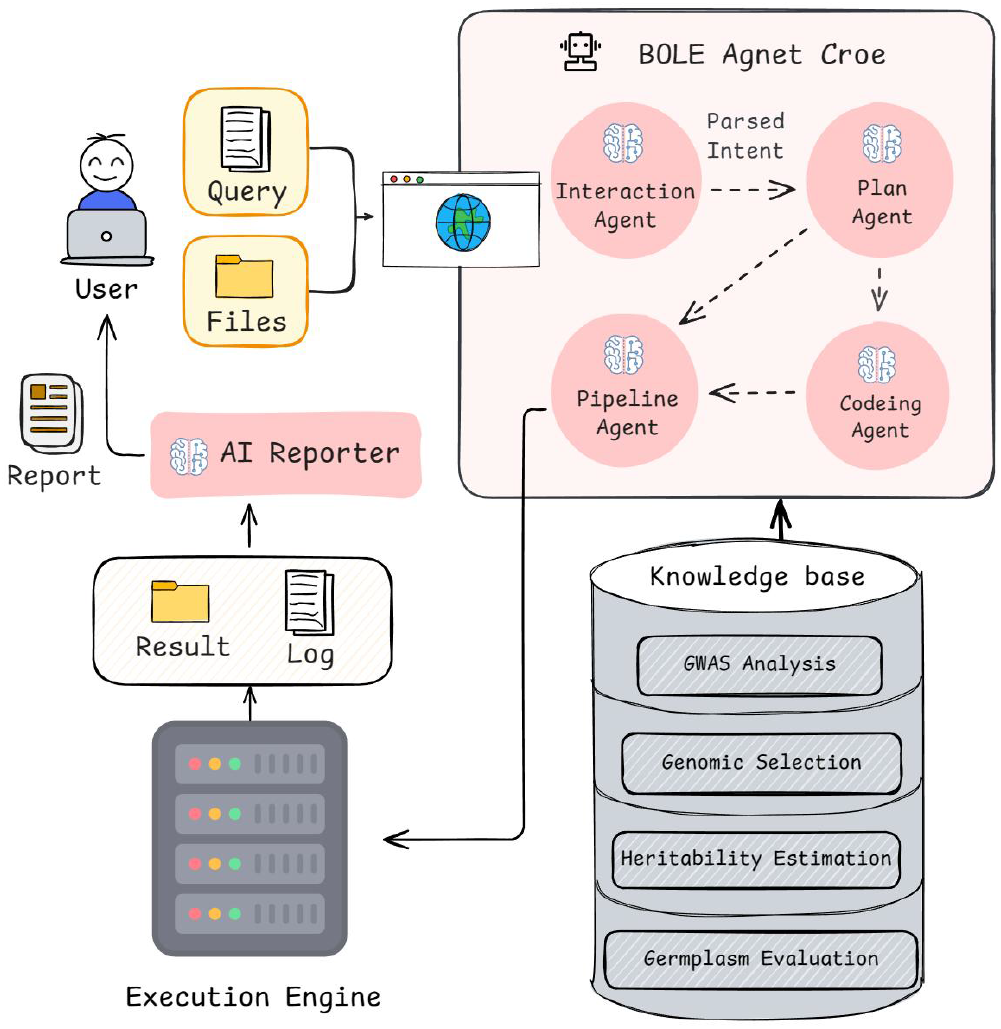
Overall architecture of the BOLE platform.

### 2.2 Integration of Large Language Models

Open-source LLMs are employed in BOLE as reasoning components to support natural language understanding and workflow planning. In the current implementation, models from the DeepSeek^18^ and Qwen families^19^ are accessed through application programming interfaces to provide semantic interpretation and decision support for individual agents. Importantly, LLMs are not used as autonomous executors of analytical logic. Instead, they operate under a knowledge-constrained multi-agent framework, where all decisions must comply with the structured script knowledge base that encodes input–output contracts, software dependencies, and domain-specific analytical rules^20^. When existing scripts cannot fully satisfy user-specified analytical requirements, a dedicated code generation agent may synthesize auxiliary scripts. Such generated scripts are constrained by predefined schemas and undergo validation before execution. This design ensures that core workflow structure and execution behavior are governed primarily by deterministic rule-based reasoning rather than unconstrained language generation. Consequently, the MAS architecture is model-agnostic and supports substitution of different LLM backends without altering workflow logic or analytical results. Reproducibility is maintained through explicit constraint checking, artifact preservation, and controlled execution^21^.

### 2.3. Formalization of the Breeding Knowledge Base

To enable autonomous reasoning, the system incorporates a structured breeding knowledge base formalized in JSON format (Figure 2). Each analytical module is represented as an atomic script unit annotated with standardized metadata, including required input formats, generated outputs, configurable parameters, software dependencies, and semantic constraints. Input–output (I/O) relationships between scripts are explicitly defined, allowing the system to reason about compatibility and automatically insert intermediate processing steps when necessary. This formalization transforms traditional bioinformatics scripts into interoperable computational building blocks, enabling autonomous assembly of complex analytical workflows.

**Figure 2.**
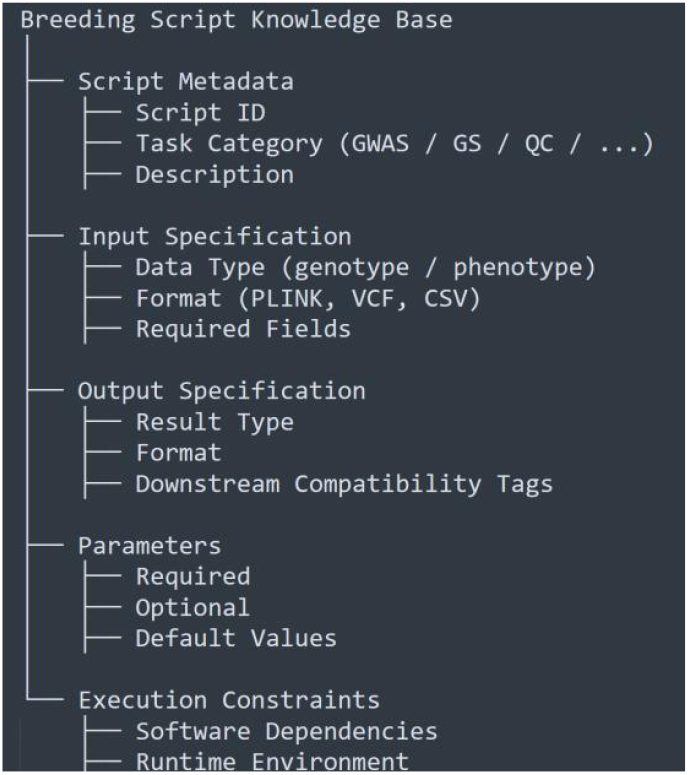
Formal structure of the breeding script knowledge base used by BOLE. Each analytical script is represented by structured metadata describing its functional role, input-output specifications, parameter constraints, and execution dependencies. This formalized representation enables agent-level reasoning over task compatibility and workflow composability, independent of any specific analytical implementation.

### 2.4. Implementation of Core Analytical Modules

The system integrates several industry-standard bioinformatics tools within its automated execution environment: Genome-Wide Association Studies: The platform supports both linear regression models via PLINK^5^ and mixed-model linear association (MLMA) via GCTA^6^. The pipelines include automated post-processing scripts to generate Manhattan and identify significant variants. Genomic Selection: Genomic prediction was performed using multiple statistical models, including BayesA, BayesB, BayesC^22^, BRR, BL^23^, and GBLUP^3^. Prediction accuracy was evaluated using K-fold cross-validation, and model performance was assessed by the correlation between predicted and observed phenotypic values. The model with the highest and most stable predictive ability was selected. Heritability Estimation:Trait heritability was estimated using linear mixed models. When only phenotypic records were available, genetic and residual variances were inferred from phenotypic data using REML^24^, and heritability was calculated as the ratio of genetic variance to total phenotypic variance. When genotype data were available, genomic heritability was estimated by fitting a mixed model with a genomic relationship matrix constructed from genome-wide SNPs^25^. Germplasm Evaluation: Population structure is assessed through unsupervised clustering using ADMIXTURE^26^. The system automates the selection of optimal ancestral populations through cross-validation error analysis and generates individual-level ancestry barplots.

### 2.5 Web Server Implementation and Availability

BOLE is deployed as a publicly accessible web-based platform that provides a lightweight interaction layer between users and the underlying multi-agent system. The web server accepts genotype and phenotype datasets together with high-level analytical intents, which are forwarded to the autonomous reasoning and execution modules. The web interface itself does not encode analytical logic or predefined workflows. All task decomposition, workflow synthesis, dependency resolution, and execution scheduling are performed exclusively by the multi-agent backend through reasoning over the script knowledge base. This separation ensures that the scientific contribution of BOLE lies in its autonomous orchestration capabilities rather than interface-level customization. The BOLE web server is freely available at http://114.80.39.6:8089/. Detailed usage instructions, example datasets, and interface demonstrations are provided in the Supplementary Materials.

## 3. Results

### 3.1 Autonomous Translation from User Intent to Executable Workflows

To evaluate the autonomous reasoning capability of BOLE, we first examined how the system translates high-level analytical intents into executable genomic breeding workflows. Instead of requiring users to manually define analysis steps, software dependencies, or execution order, BOLE accepts abstract task descriptions and autonomously constructs complete workflows through multi-agent coordination. Upon receiving a user intent, the planning agent decomposes the request into atomic analytical objectives and queries the script knowledge base to identify compatible modules. Input–output constraints, software dependencies, and parameter requirements are resolved dynamically, allowing the system to infer intermediate processing steps such as genotype quality control, format harmonization, and data validation (Figure 3 & Figures S1-3). This process enables BOLE to generate executable workflows without predefined static pipelines, demonstrating its ability to reason over heterogeneous tools and adapt analysis trajectories to varying data conditions and user intents.

**Figure 3.**
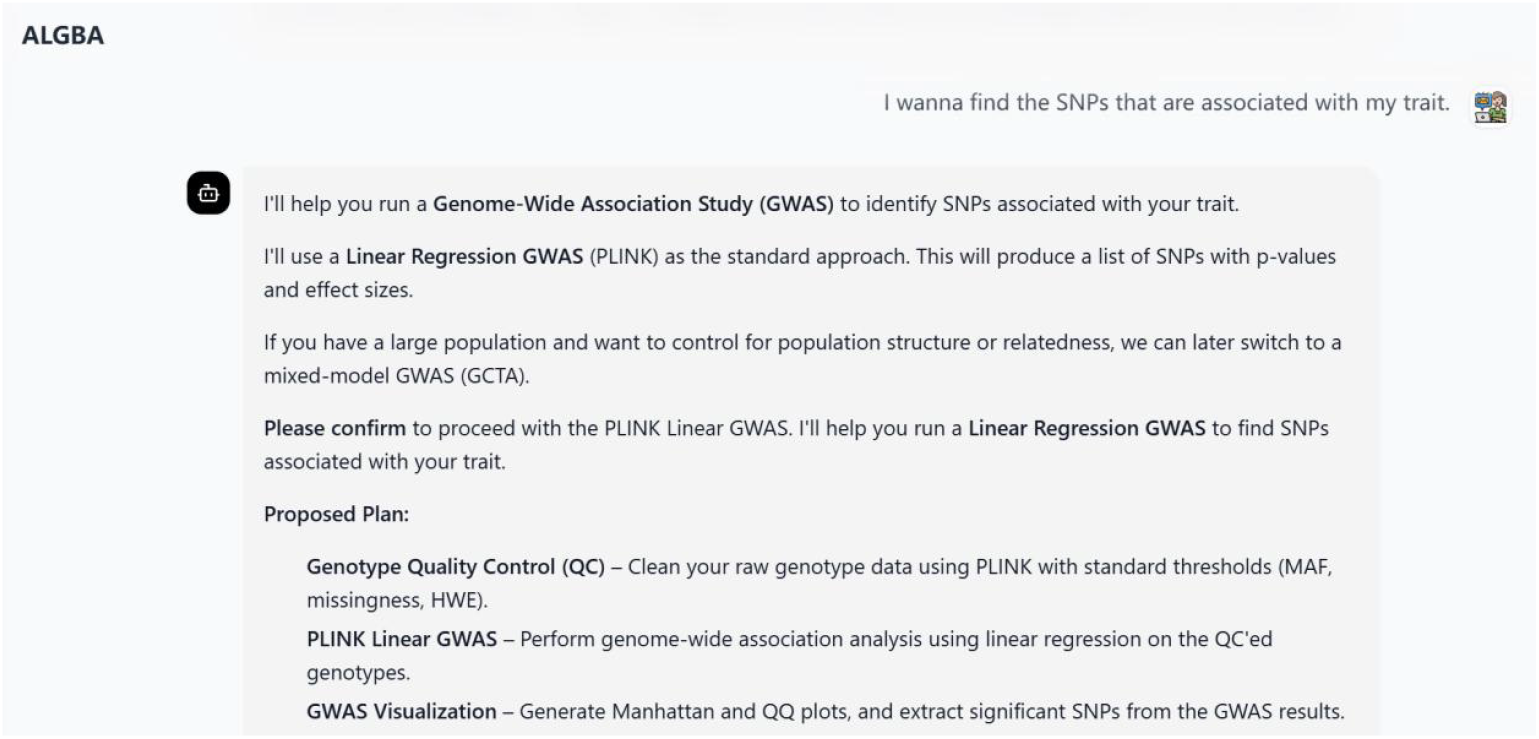
Example of an autonomously generated genomic breeding workflow for GWAS analysis. Given a high-level analytical intent, BOLE dynamically decomposes the task into atomic analysis modules and constructs an executable workflow.

### 3.2 End-to-End Genomic Breeding Analysis Across Core Functional Modules

We next assessed BOLE’s ability to perform end-to-end genomic breeding analyses spanning its four core functional modules: GWAS, heritability estimation, germplasm evaluation, and GS. Starting from raw genotype and phenotype datasets, the system autonomously executed all required preprocessing, statistical modeling, and result aggregation steps. Heritability estimates were computed using genomic relationship matrices, providing quantitative assessments of genetic contribution (Figure 4A). Germplasm evaluation analysis produced ADMIXTURE-based population structure outputs. For genomic selection, BOLE coordinated multiple prediction models and performed internal model evaluation, producing predictive accuracy metrics under standardized cross-validation settings (Figure 4C). GWAS analyses generated Manhattan (Figure 4D), enabling the identification of loci associated with target traits. All analytical outputs were generated within a single autonomous workflow, without manual intervention or reconfiguration between modules. Together, these results demonstrate that BOLE can seamlessly integrate multiple breeding-relevant analyses into a coherent, end-to-end pipeline, reflecting real-world genomic breeding scenario.

**Figure 4.**
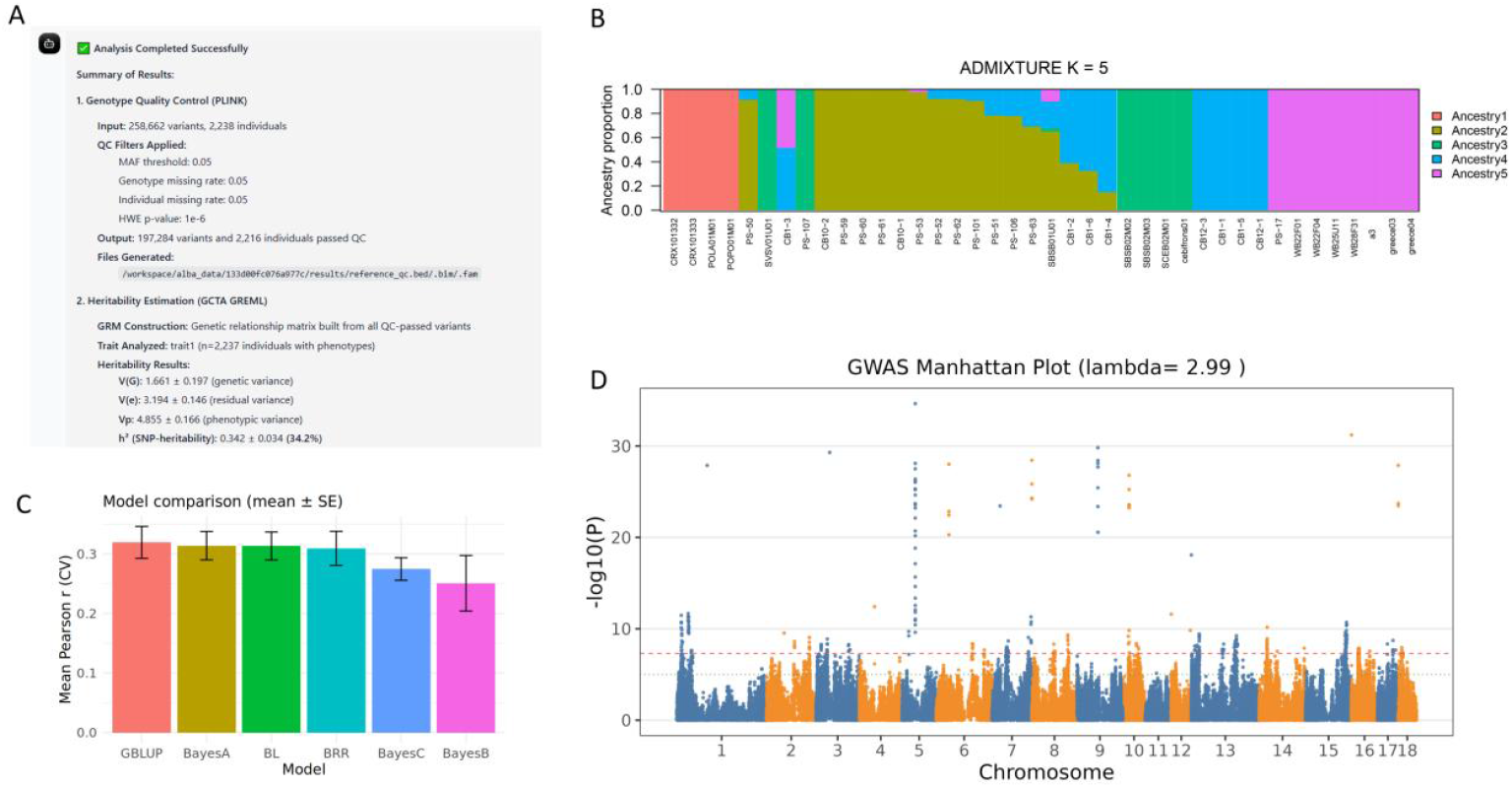
Representative outputs generated by BOLE across core genomic breeding modules. BOLE autonomously performs heritability estimation (A), Germplasm evaluation (B), genomic selection with model evaluation (C), and genome-wide association studies (D) within a unified workflow. All results are generated from raw genotype and phenotype inputs without manual reconfiguration between analytical steps, demonstrating the platform’s end-to-end breeding analysis capability.

### 3.3 Consistency and Reproducibility of Autonomous Workflows

Reproducibility is a critical requirement for genomic data analysis systems. To assess the consistency of BOLE’s autonomous execution, identical analytical intents and datasets were submitted in repeated runs. Across independent executions, the system consistently generated identical workflow structures. The deterministic reasoning over the script knowledge base ensured that identical input conditions yielded identical workflow (Figure S4), while execution artifacts—including intermediate files and final outputs—were preserved and traceable (Figure 5). This artifact-aware execution model enables transparent inspection of each analytical step and facilitates reproducible research practices. These observations indicate that BOLE maintains reproducibility despite its dynamic, intent-driven workflow generation, addressing a common concern associated with automated analysis systems.

**Figure 5.**
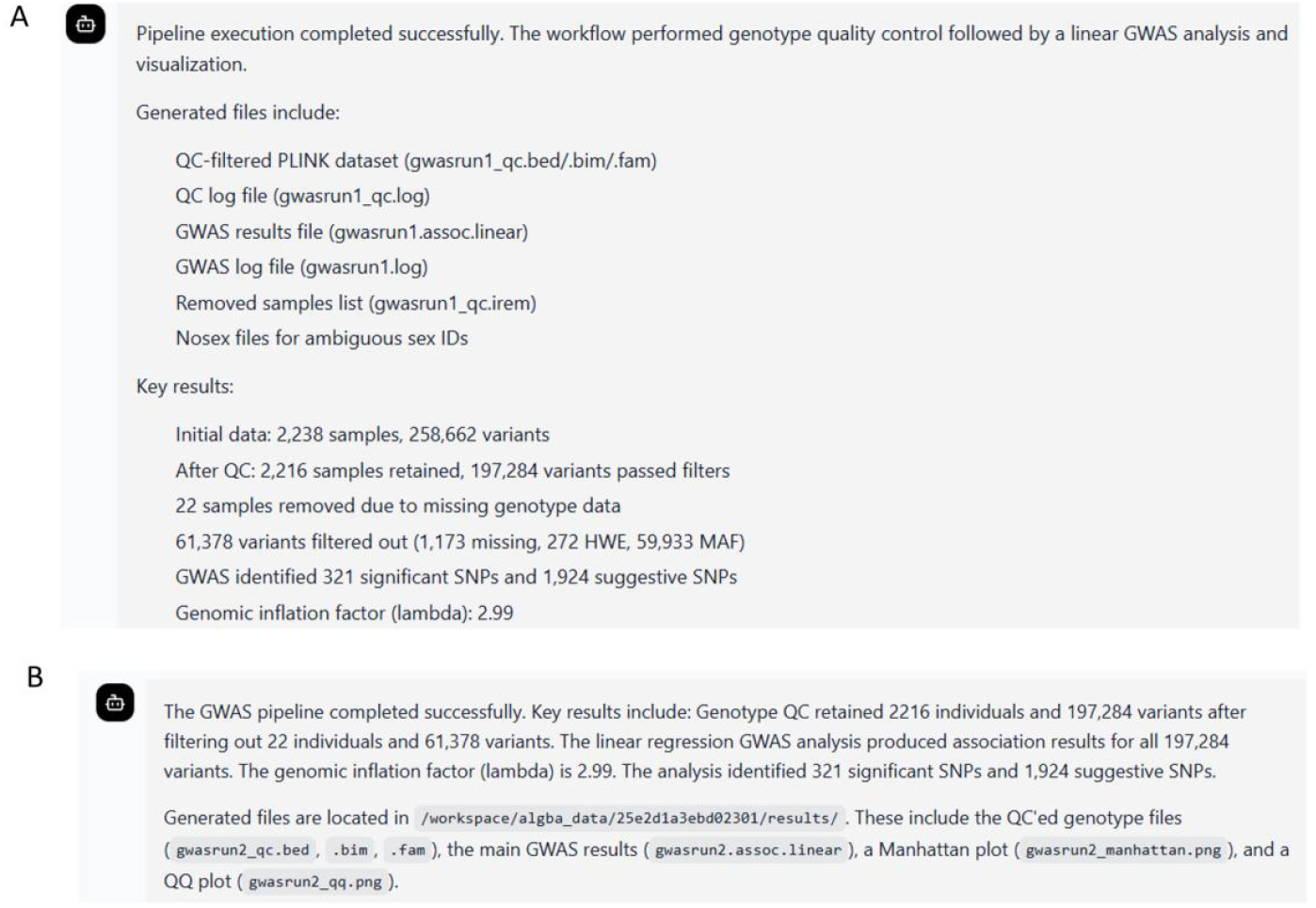
Reproducibility of autonomously generated workflows. Repeated executions of identical analytical intents and datasets result in consistent workflow structures, parameter configurations, and analytical outputs. The deterministic reasoning over the script knowledge base ensures reproducible execution despite the dynamic, intent-driven nature of workflow generation.

### 3.4 Dynamic Cross-Task Workflow Composition

To evaluate BOLE’s ability to perform non-fixed, intent-driven analytical workflows, we designed a multi-stage breeding analysis scenario in which downstream tasks explicitly depended on intermediate results generated in prior analyses. In this scenario, a user first requested a GWAS for a target trait and subsequently instructed the system to perform GS based on the GWAS results. Upon receiving the initial GWAS request, BOLE autonomously constructed and executed a complete association analysis workflow, including genotype quality control, phenotype alignment, and statistical association testing. The resulting intermediate artifacts, such as association statistics and selected marker sets, were recorded and indexed within the system’s execution context. The autonomously generated analytical plan is the same as figure 3, where the system dynamically composes a multi-stage workflow without relying on predefined composite pipelines.

When the subsequent GS request was issued, BOLE reasoned over the available intermediate artifacts and identified compatible input-output relationships for downstream genomic prediction. Rather than triggering a fixed GS pipeline, the system verified data compatibility, selected valid parameter configurations, and assembled an executable genomic selection workflow that incorporated the upstream GWAS outputs (Figure 6A). The execution trace of this dynamically constructed workflow, including module-level execution status and preserved artifacts, is shown in Figure 6B. Together, these results demonstrate that BOLE supports dynamic cross-task reasoning and execution beyond isolated, task-specific workflows. By enabling adaptive composition of multi-stage analytical processes based on user intent and intermediate results, BOLE moves beyond static workflow automation and provides a flexible framework for complex breeding analysis scenarios commonly encountered in real-world genomic breeding programs.

**Figure 6.**
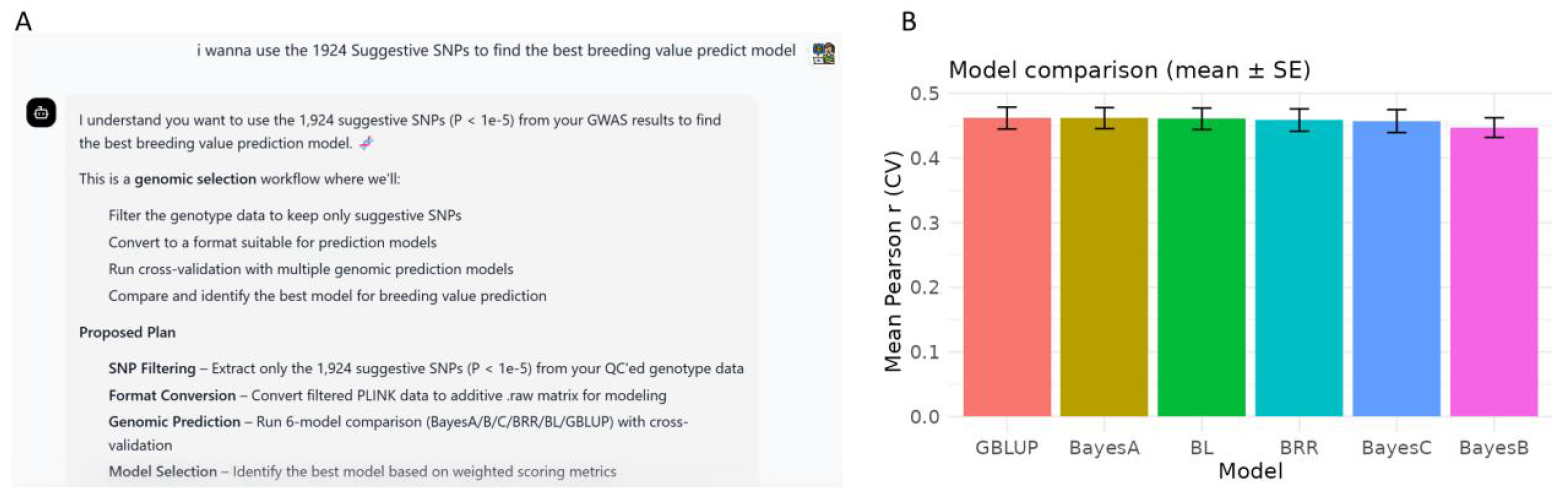
Autonomous cross-task workflow planning and execution in BOLE. (A) Illustration of an adaptive analytical plan generated by BOLE in response to a new user requirement after GWAS analysis. The system autonomously integrates GWAS results into subsequent genomic breeding analyses and formulates a coherent multi-step workflow. (B) Results produced by executing the adaptive workflow, demonstrating BOLE’s capability to support flexible, cross-task genomic breeding analyses within a unified framework.

## 4. Discussion

The increasing complexity of genomic breeding analyses has created a substantial gap between methodological advances and their practical adoption by breeding researchers. While numerous statistical models and software tools have been developed for genomic breeding, their effective use still requires extensive computational expertise and manual workflow engineering^27^. BOLE addresses this gap by introducing an autonomous, intent-driven analysis framework that emphasizes reasoning, workflow composition, and execution traceability rather than the invention of new analytical models. A central contribution of BOLE lies in its departure from predefined, static pipelines. Conventional workflow systems require users to anticipate analytical sequences in advance and encode them explicitly, which limits flexibility when analytical goals evolve or intermediate results suggest new directions^28^. In contrast, BOLE dynamically synthesizes workflows based on user intent and the compatibility constraints defined in a structured script knowledge base. As demonstrated in the Results, identical analytical intents consistently produce identical workflow structures and execution artifacts, indicating that autonomy in workflow generation does not compromise reproducibility. This paradigm shift — from fixed pipelines to intent-driven orchestration—represents a meaningful advance in the usability of genomic breeding analyses.

The integration of large language models (LLMs) within BOLE may raise concerns regarding stochastic behavior and reproducibility^29^. Importantly, LLMs in BOLE do not function as unconstrained generators of analytical logic, but rather as constrained reasoning components operating within explicitly defined system boundaries^30^. Workflow planning and execution must satisfy deterministic input-output specifications, software dependencies, and artifact compatibility rules encoded in the script knowledge base. Even in cases where the system generates auxiliary scripts to bridge incompatible formats or unmet analytical requirements, such code generation is guided by strict functional constraints and is fully recorded as execution artifacts. This design ensures that analytical outcomes are governed by system-level determinism rather than free-form language generation, thereby preserving reproducibility and transparency^31^.

Another notable feature of BOLE is its ability to support non-fixed, cross-task analytical workflows. In practical breeding scenarios, analyses are rarely isolated; results from one task, such as GWAS, often motivate downstream analyses including genomic selection or candidate evaluation^32^. While such multi-stage workflows can be manually designed by experienced analysts, these solutions are typically ad hoc, fragile, and difficult to generalize. BOLE demonstrates the capacity to autonomously reason over intermediate outputs and assemble compatible downstream workflows without predefined composite pipelines. This capability highlights the system’s adaptability and its potential to support exploratory and iterative analytical processes common in real-world breeding programs.

Beyond automation, BOLE emphasizes artifact-aware execution and traceability. Every analytical step, including intermediate files, parameters, execution logs, and dynamically generated scripts, is preserved and accessible for inspection. This design not only supports reproducible research practices but also facilitates methodological transparency, enabling users to understand, audit, and reuse analytical results. Such traceability is particularly important in breeding research, where analytical decisions can have long-term biological and economic implications.

Despite these strengths, several limitations warrant discussion. BOLE does not aim to replace expert knowledge in breeding or statistical genetics, nor does it propose novel analytical models. Instead, it serves as an enabling infrastructure that lowers the technical barriers to applying established methods. The system’s performance and analytical breadth are also constrained by the completeness and quality of the script knowledge base; extending BOLE to new analytical domains requires careful curation of compatible scripts and metadata. Furthermore, while the current implementation demonstrates robustness across core genomic breeding tasks, large-scale deployment and broader community adoption will require continued validation and maintenance^21^.

In summary, BOLE represents a step toward intelligent, autonomous genomic breeding analysis by reframing workflow construction as a reasoning problem rather than a manual engineering task. By combining intent-driven planning, knowledge-constrained autonomy, and artifact-level reproducibility, the system bridges the gap between methodological innovation and practical usability. This framework provides a foundation for future developments in intelligent bioinformatics systems and offers a scalable path toward more accessible and reproducible genomic breeding analyses.

## Acknowledgements

This work was supported by grants from the National Key R&D Program of China (Grant NO. 2021YFD301201, 2024YFF0728800), Special Funding for the Construction of the High-Level Academy of Agricultural Sciences (No. NYQS202632) and the Shenzhen Outstanding Talents Training Fund (Grand NO. 202102).

## Author contributions

This study was conceived by K.L. and Z.W.. The platform was constructed by Z.W. and X.L.. Z.W. wrote the manuscript. All authors have read and approved this manuscript.

