## Supplementary figures for "BOLE: A Knowledge-Enhanced Multi-Agent Framework for Intelligent Genomic Breeding"

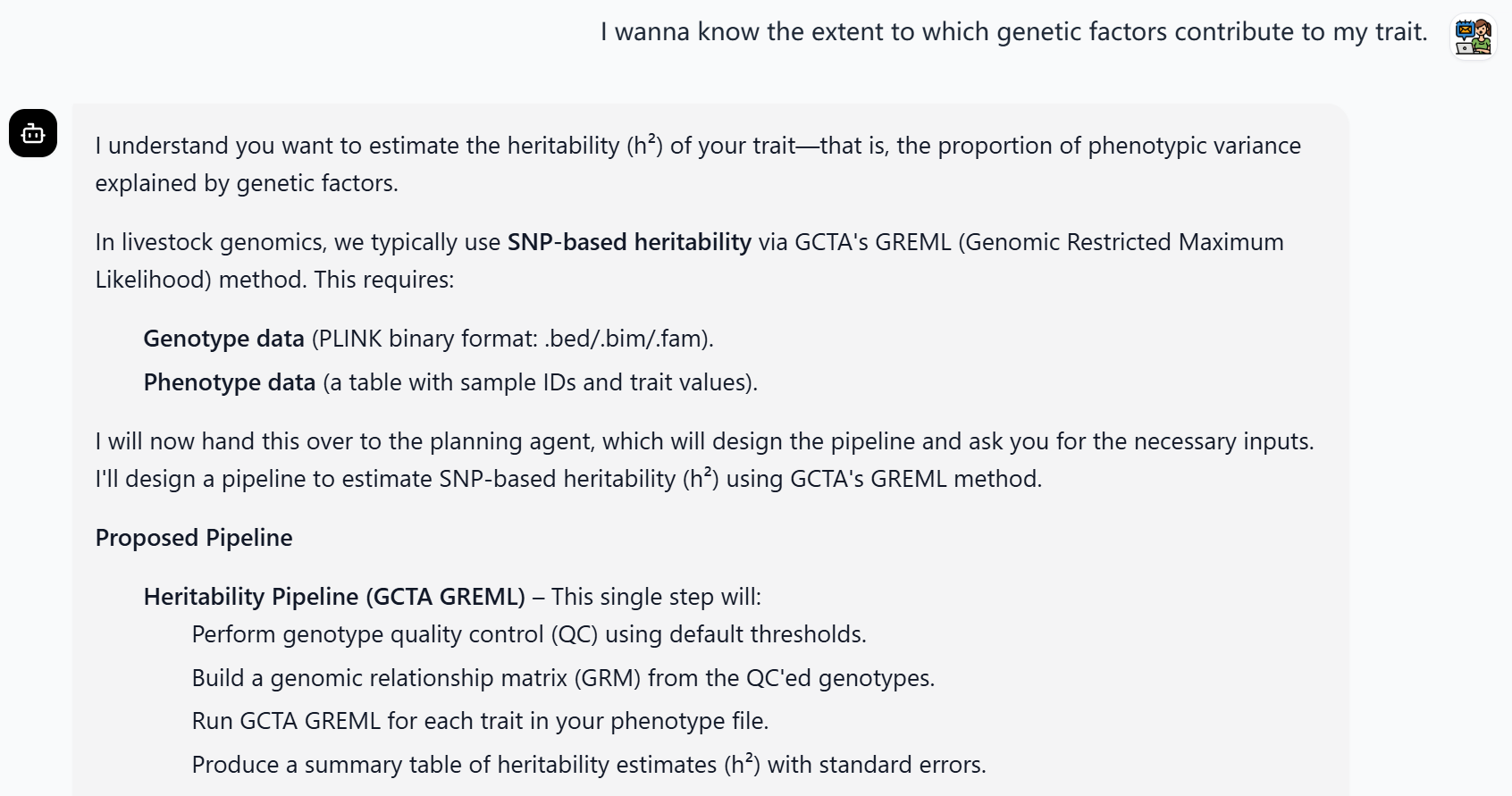
**Figure S1. Autonomous generation of a** heritability estimation **analysis workflow in BOLE.**

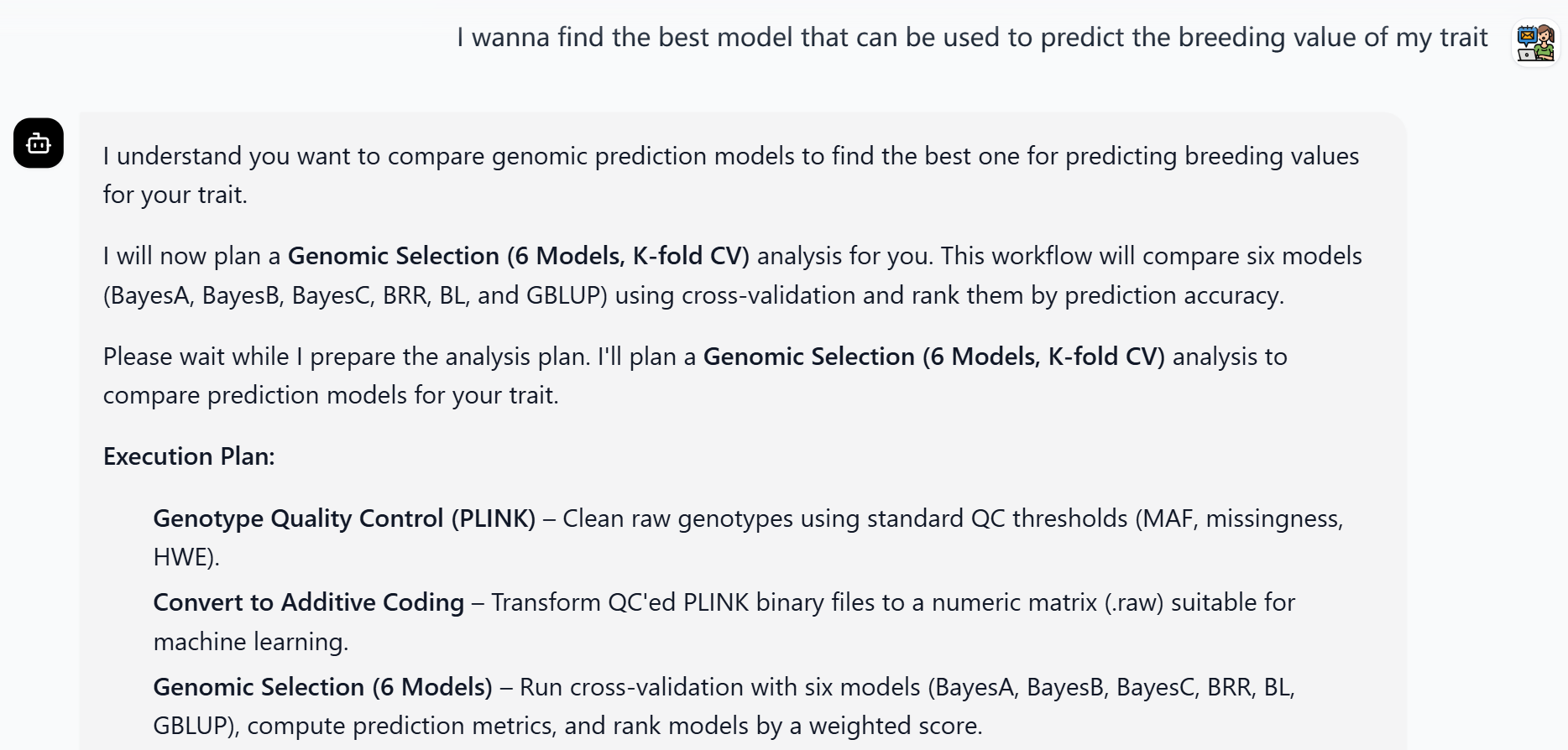
**Figure S2. Autonomous generation of a genomic selection pipeline in BOLE.**

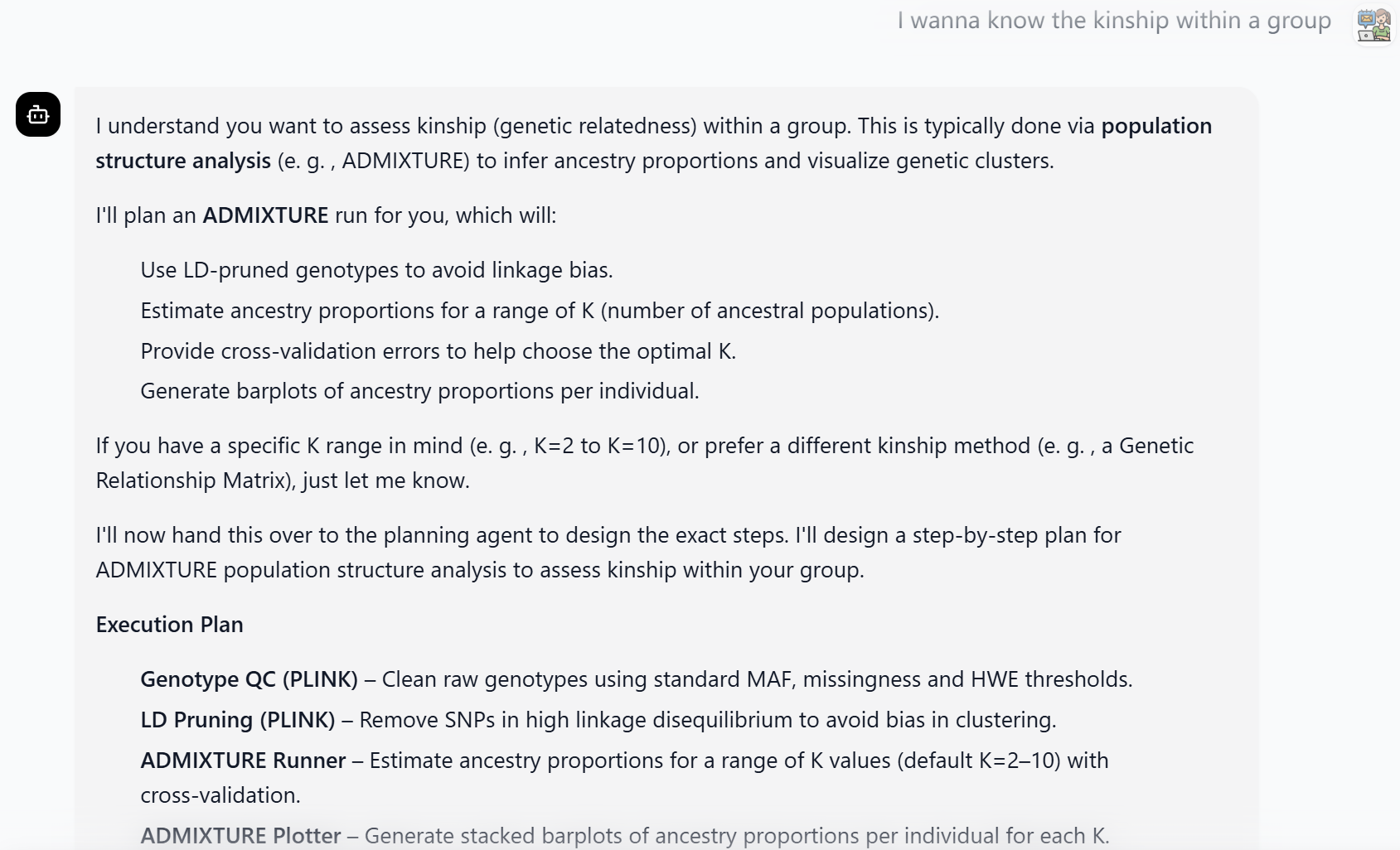
**Figure S3. Autonomous generation of a germplasm evaluation pipeline in BOLE.**

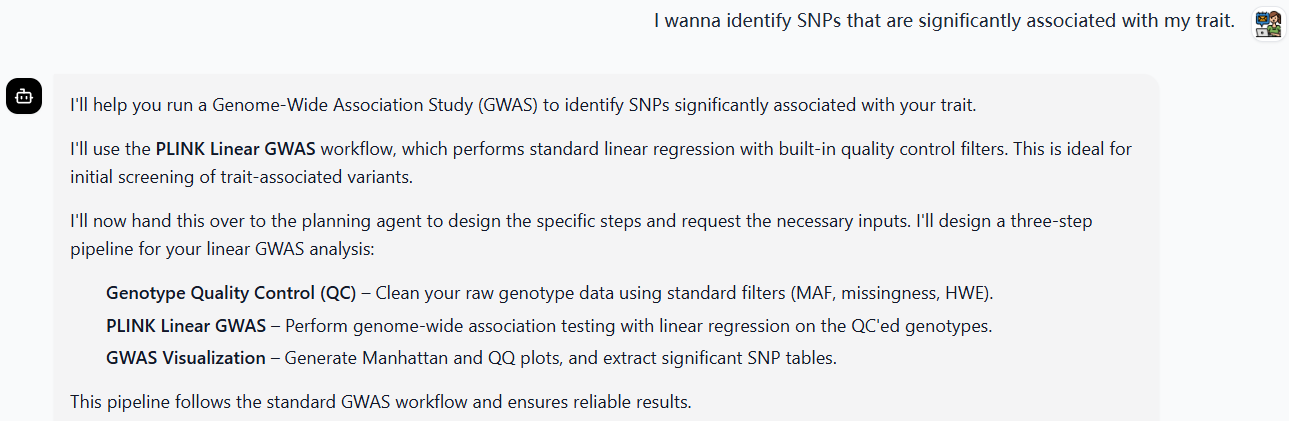

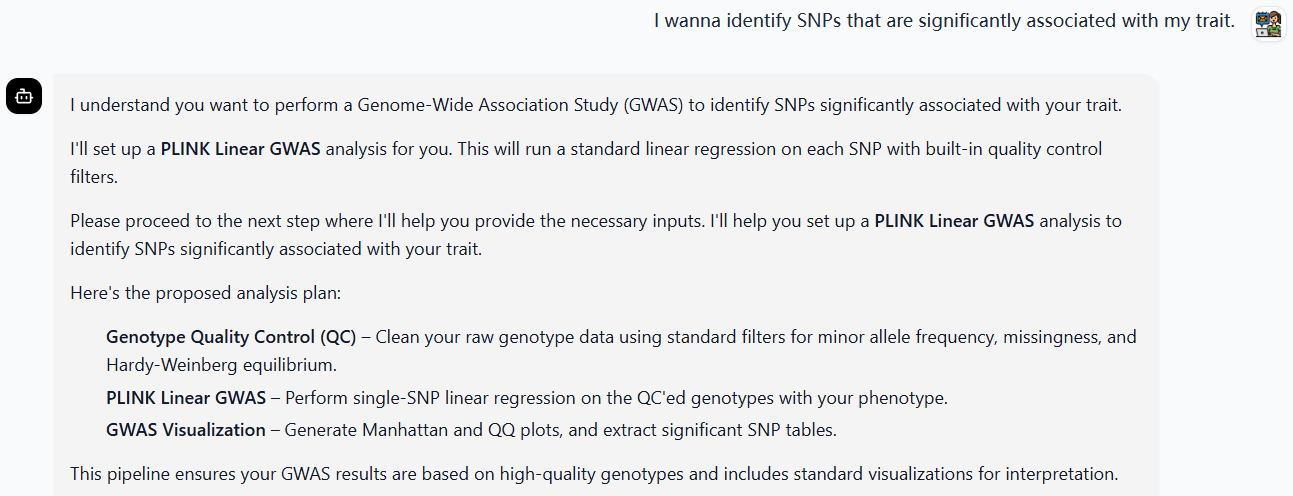
**Figure S4. Consistency of autonomously generated workflow plans across independent executions.** Two independent executions of the same analytical intent and dataset produced identical workflow structures. The sequence of analytical modules and their dependencies remained unchanged between runs, demonstrating that BOLE generates stable and deterministic workflow plans despite its intent-driven reasoning mechanism.
